# Drastic shift in flowering phenology, an instant reproductive isolation mechanism, explains the population structure of *Imperata cylindrica* in Japan

**DOI:** 10.1101/2020.06.30.179440

**Authors:** Yasuyuki Nomura, Yoshiko Shimono, Nobuyuki Mizuno, Ikuya Miyoshi, Satoshi Iwakami, Kazuhiro Sato, Tohru Tominaga

**Affiliations:** Research Institute for Food and Agriculture, Ryukoku University, 1-5 Yokoya, Seta Oe-cho, Otsu, Shiga 520-2194, Japan; Graduate School of Agriculture, Kyoto University, Kitashirakawa Oiwake-cho, Sakyo, Kyoto 606-8502, Japan; Institute of Plant Science and Resources, Okayama University, 2-20-1 Chuo, Kurashiki, Okayama 710-0046, Japan

**Keywords:** hybridisation, flowering phenology shift, ecotype, instant reproductive isolation, *Imperata cylindrica*, population genetic structure

## Abstract

Reproductive isolation plays an important role in population differentiation and speciation, thus enhancing biodiversity in wild plants. Hybridisation sometimes involves rapid reproductive isolation between parents and their hybrids through the novel traits of hybrids derived from a new combination of genomes. Here, we report how a hybrids’ new phenotype contributes to rapid reproductive isolation between two ecotypes of *Imperata cylindrica*. The two ecotypes differ in their flowering phenology and habitats. An analysis with genetic markers revealed that hybrid populations consisted of only F_1_ individuals. Both parental ecotypes flowered in spring, but F_1_s flowered in fall. This drastic shift in flowering phenology prevented backcrossing parental ecotypes to F_1_. F_1_s flowered in fall and dispersed seeds in winter. The germination percentage of seeds set on F_1_ was extremely low in their habitats, and seedlings did not survive due to the low temperatures in winter, resulting in the absence of a F_2_ generation. In conclusion, flowering phenology mismatch promotes reproductive isolation between parents and F_1_, resulting in a hybrid population consisting of only F_1_s.

## Introduction

Around 50% of flowering plants are estimated to have experienced polyploid speciation. Nearly all such polyploid individuals are considered to be derived from hybridization (Soltis and Soltis 2009; Wood et al. 2009; Soltis et al. 2015). Hybridisation is a major source of phenotypic variation (Grant and Grant 1994) and a driving force of evolution (Coyne and Orr 2004). One of the evolutional consequences of hybridisation is hybrid speciation, which cause species diversification (Mallet 2007; Soltis and Soltis 2009; Wood et al. 2009). The resultant hybrids are sometimes prevented from crossing with their parental lineages due to an altered trait related to reproduction, resulting in the change of population structure and ultimately species diversification (Buerkle et al. 2000; Duenez-Guzman et al. 2009; Kagawa and Takimoto 2018).

Previous studies have demonstrated that novel traits derived from hybridization directly contribute to reproductive isolation. Mating between individuals that have evolved independently results in a novel combination of genomes or gene sets; this process can generate a wide range of phenotypes, including some of which that exceed or interpose variations among the parental lineages by overdominance or transgressive phenotype (Johansen-Morris and Latta 2006). For example, floral color in *Iris* species (Taylor et al. 2013) and floral odor for *Narcissus* species (Marques et al. 2016) differ among hybrids from those of their parents, resulting in recruitment of novel pollinators for hybrids and reproductive isolation between hybrids and parents. Similarly, new hybrid phenotypes related to beak morphology and mating songs in Darwin’s finch (Lamichhaney et al. 2018), wing color patterns in *Heliconius* butterfly (Mavárez et al. 2006), and behavioral mate choice in cichlids (Selz et al. 2014) have caused reproductive isolations from parents. Thus, novel hybrid traits can generate genetic and evolutionary changes.

In this study, we report a novel trait in hybrids of two *Imperata cylindrica* (L.) Raeusch (cogongrass) ecotypes. *Imperata cylindrica* is a perennial rhizomatous grass with a self-incompatible and wind-pollinated reproduction system; it is native to tropical and subtropical areas of the Northern and Southern Hemispheres, including Japan (Holm et al. 1977). Japanese cogongrass populations consist of two ecotypes: common type (C-type) and early flowering type (E-type) (Matumura and Yukimura 1980; Tominaga et al. 1989; Mizuguti et al. 2003). These ecotypes are typically distinguished by their morphology; the E-type has a glabrous culm, whereas the C-type has a hairy culm (Figure 1). They also differ in terms of habitat and flowering phenology: the C-type mainly lives in dry habitats (*e*.*g*. roadside, levee of paddy fields), whereas the E-type often lives in wet habitats (*e*.*g*. marshy area, moist fallow fields). In addition, the E-type flowers approximately 1 month earlier than the C-type. These differences in habitat and flowering phenology isolate the two ecotypes, although the existence of hybrids between the two ecotypes was initially recognized through allozyme analysis using a single marker (Mizuguti et al. 2004; Tominaga et al. 2007). Since then, the distribution and genetic population structure of the hybrids remained unknown.

**Figure 1.**
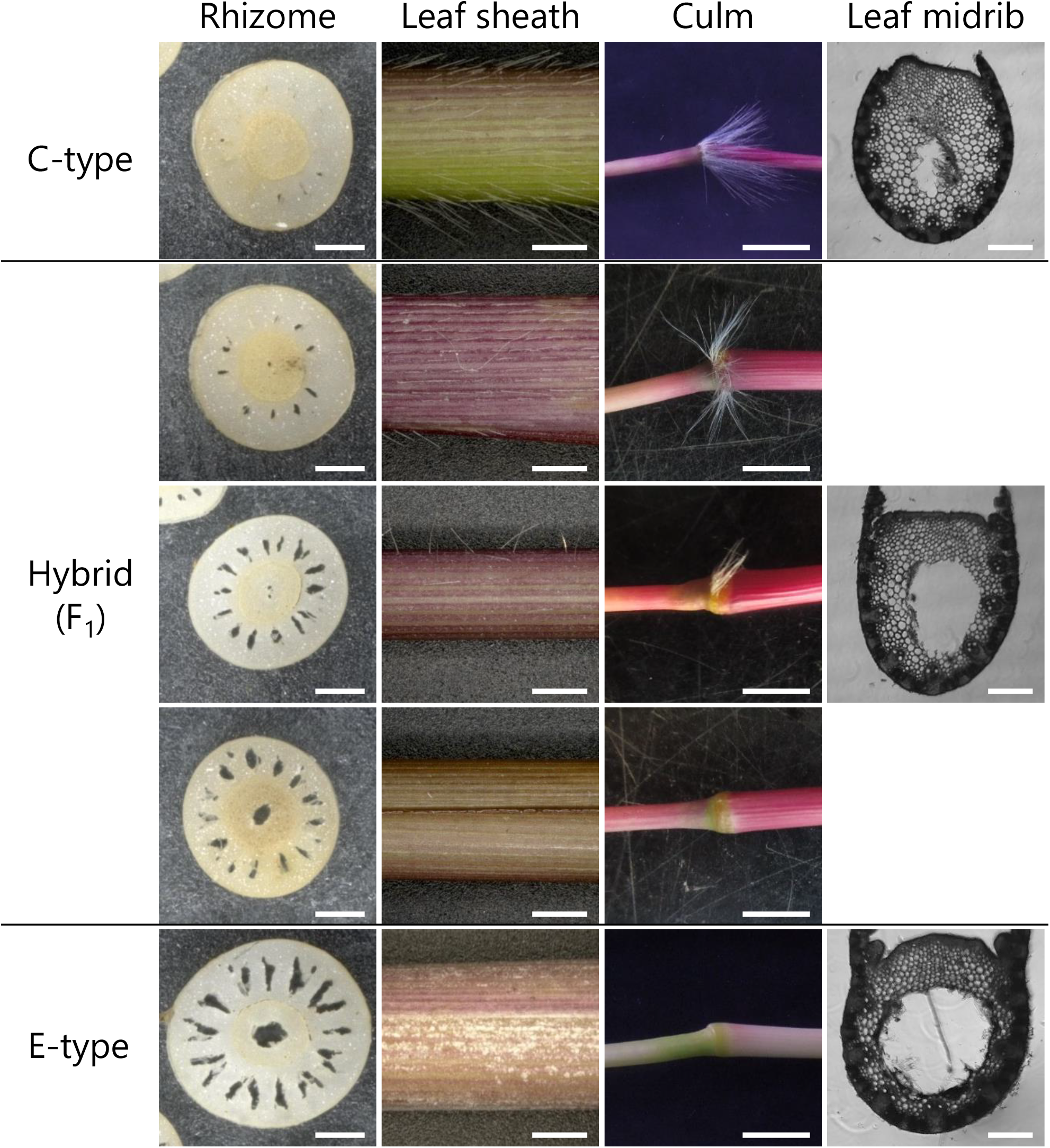
Morphology of C-, E-type and hybrids. Bars in right bottom of photos are scale bars. Scale bars mean 1 cm, 1 cm, 0.5 cm and 250 μm in rhizomes, leaf sheathes, culms and leaf midribs, respectively.

In this study, we analysed 183 populations of cogongrass collected from the 1980s to the 2010s throughout Japan using newly established molecular markers. We performed field surveys and common garden experiments to determine the outcomes of backcrossing and crossing among the F1 generation to evaluate the contribution of hybrid flowering phenology to their genetic population structure. Our findings will provide insight into the reproductive isolation resulting from hybridisation.

## Materials and methods

### Accession and study site

Three hundred fifty accessions of cogongrass were collected from throughout Japan, from 1980s to 2010s, and maintained at an experimental farm of Kyoto University, Kyoto, Japan (35°01′54.5″N, 135°46′59.5″E) (Figure 2; Table S1). The chloroplast DNA (cpDNA) haplotypes of 33 out of the 350 accessions were previously determined by Nomura et al. (2015). Field survey and material collection were conducted in north sites (Aomori, Akita, Iwate, Yamagata, Miyagi, Fukushima and Tochigi Prefecture; 36°54′41″N – 40°32′07″N, 139°48′34″E – 141°40′10″E) and south sites (Wakayama Prefecture; 33°29′01″N–33°39′39″N, 135°46′39″– 135°57′59″) regions, Japan in 2016–2018 (Figure S1).

**Figure 2.**
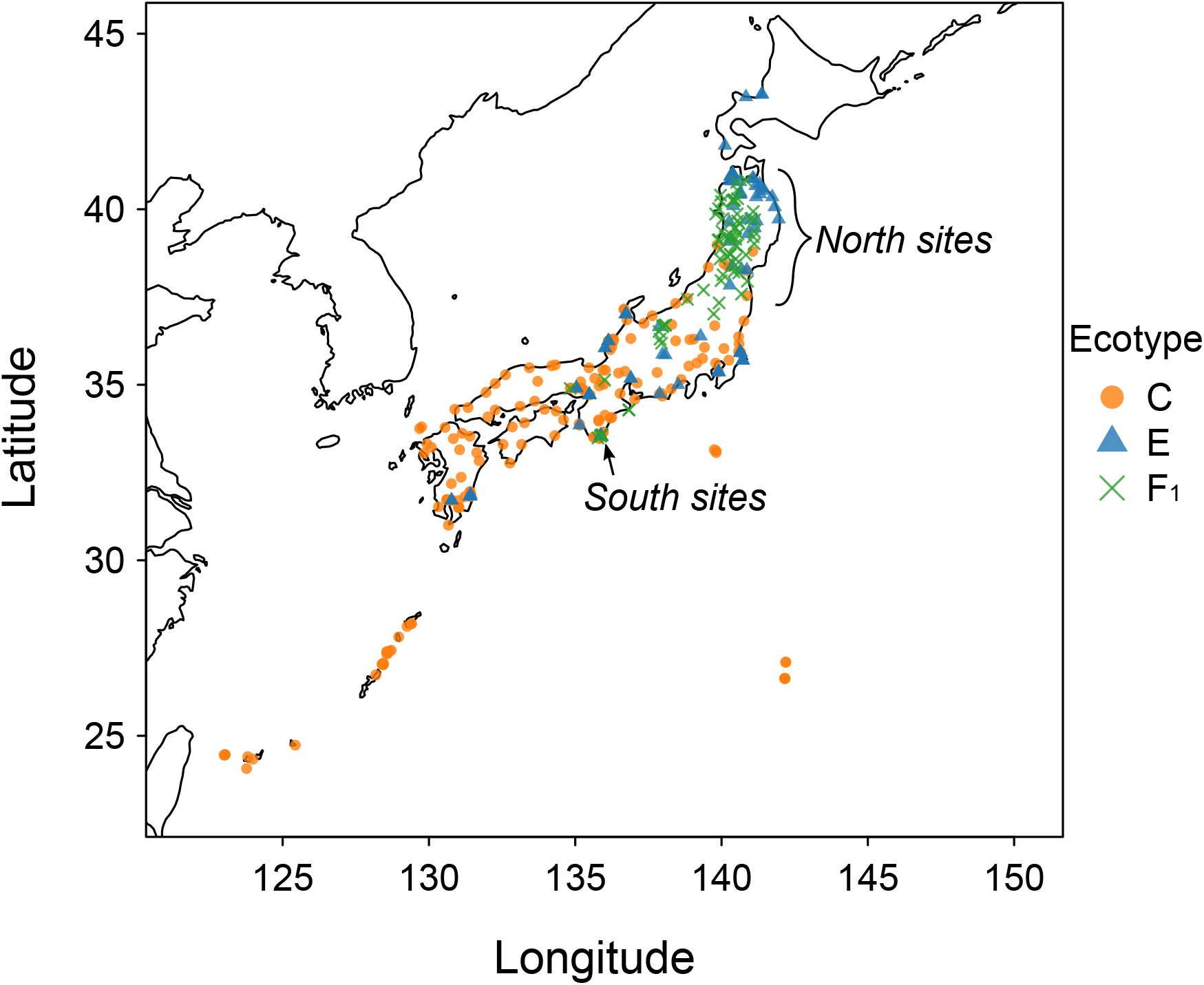
Sampling location and distribution of each ecotype. Ecotypes were determined using CAPS marker in ITS region.

### Evaluation of morphological traits

Morphological traits were evaluated for a total of 46 accessions: 17 accessions of the C-type, 15 accessions of the E-type, and 14 accessions of putative hybrids (Table S1). The existence of hairs on culm nodes (1, absence; 2, partial presence; 3, presence), hairs on leaf sheathes (1, absence; 2, partial presence; 3, presence), wax on leaf sheathes (1, absence; 2, presence), the ratio of aerenchyma diameter to midrib diameter, and the ratio of aerenchyma diameter in a pith to rhizome diameter were recorded (Figure 1).

### Nucleic acid extraction

DNA was extracted from leaves of cogongrass using the modified cetyltrimethylammonium bromide method (Murray and Thompson 1980). Total RNA was extracted from leaves using a RNeasy Plant Mini kit (Qiagen, Tokyo, Japan) according to the manufacturer’s instructions.

### Development of nuclear markers

Total RNA extracted from 22 accessions of the E-type throughout Japan (Table S1) was bulked, and a total of 4 μg of total RNA was used to construct paired-end libraries using the TruSeq RNA Sample Preparation Kit v3 (Illumina, Inc., San Diego, CA, USA) according to the manufacturer’s instructions. The libraries were sequenced on MiSeq (Illumina) with 300-bp paired-end reads. Filtering of low-quality bases (Phred score < 20) and adapter sequences, *de novo* transcriptome assembly, and detection of polymorphisms were conducted using CLC Genomics Workbench software version 8.0 (CLC bio Japan, Inc., Tokyo, Japan).

One hundred seven primer sets were designed to amplify DNA of 300–900 bp in which polymorphisms were detected in the above RNA-Seq analysis. PCR was conducted using DNA from two C-type and six E-type accessions selected at random. PCR amplification was performed in a 20-μL volume containing 2.5–25 ng of extracted DNA, 0.2 mM each dNTP, 0.5 μM each forward and reverse primer, 2.0 μL of 10× ThermoPol™ Buffer, and 0.5 U of *Taq* DNA polymerase (New England Biolabs Japan Inc., Tokyo, Japan). The PCR programme consisted of an initial denaturation at 95.0°C for 3 min followed by 45 cycles at 95.0°C for 15 s, annealing 55°C for 30 s, and extension at 68.0°C for 1 min; with a final extension at 68.0°C for 3 min. The PCR products were subjected to agarose gel electrophoresis. The PCR products that contained a single band of the expected size were purified using ExoSAP-IT (USB Corporation, Cleveland, OH, USA) and directly sequenced using an ABI 3130xl Genetic Analyzer (Applied Biosystems, Foster City, CA, USA) with BigDye™ Terminator v. 3.1 (Applied Biosystems). Fifty seven PCR products were sequenced. Among the 57 primer sets that resulted in PCR products with a clear chromatogram in direct sequencing, 10 were selected for investigation of population structure (Table S2).

In addition to the 10 primer sets, 2 primer sets were designed based on the sequences of cogongrass in GenBank: *ppc-C4* (AM690231) and the ITS region (JN407507). The primers are listed in Table S2.

### Development of CAPS markers to distinguish ecotypes

Cleaved amplified polymorphic sequences (CAPS) markers in nuclear DNA were developed to distinguish ecotypes and their hybrids. Primers were designed for the ITS region and EST104 (Table S2, ITS_CAPS and EST104_CAPS), and PCR was performed as described above. The PCR products of ITS and EST104 were digested with *Dde*I and *Rsa*I (New England Biolabs Japan, Inc., Tokyo, Japan), respectively, and resolved on an agarose gel (Figure S2).

A CAPS marker on cpDNA was also developed to distinguish maternal ecotypes. The *psbA–matK* region was amplified by PCR using the primer set (Yasuda and Shibayama 2006): forward PSA-F, 5′-CAGGCTTGTACTTTCGCGTC-3′; reverse MTK-R, 5′-CAATGTCCTGGTGAATCCTC-3′. PCR was carried out as described (Nomura et al. 2015). The PCR products were incubated with *Dra*I (TaKaRa Bio, Inc., Shiga, Japan) and subjected to gel electrophoresis.

### Analysis of population structure

SNPs with a low frequency (<10% in all accessions) were excluded from haplotype analysis using PHASE 2.1 (Stephens et al. 2001). Principal coordinate analysis (PCoA) was conducted using GENALEX 6.502 (Peakall and Smouse 2006, 2012). The population genetic structure was inferred using the Markov chain Monte Carlo (MCMC) and the Bayesian clustering algorithms implemented in STRUCTURE v. 2.3.1 (Pritchard et al. 2000) with 1,000,000 MCMC steps following 100,000 burn-in MCMC steps. The number of clusters (*K*) was tested from 1 to 10 with 10 replicates. The appropriate number of clusters was estimated using the Evanno *ΔK* parameter (Evanno et al. 2005).

Distinction of hybrids from parental ecotypes and identification of generation of hybrids were conducted using NEWHYBRIDS v. 1.1 (Anderson and Thompson 2002). Ten independent runs were conducted with 1,000,000 MCMC steps following 100,000 burn-in MCMC steps, and the posterior probability was computed for each of the six classes: the two-parental ecotypes, F_1_ and F_2_ generations, and backcrosses to each parental class. The 10 independent results of STRUCTURE and NEWHYBRIDS were amalgamated using CLUMPP v. 1.1.2 (Jakobsson and Rosenberg 2007).

### Genotyping of putative F_2_s

Seven F_1_ plants were open-pollinated in 15 m × 15 m plot of an experimental farm of Kyoto when only F_1_ plants were in bloom. The seeds collected from the plants were germinated on filter paper in a Petri dish at 30/20°C day/night with a 12-h photoperiod in an incubator and seedlings were transplanted into plastic pots (11.3-cm diameter × 14.0-cm height). The genotype of each putative F_2_ was evaluated using the CAPS marker in the ITS region and EST104, and direct sequencing using ppc-C4 and EST72 primer sets (Table S2) described above.

### Percentage of hybridisation between the C- and E-types in natural habitats

Seeds were collected during May to July in 2017 and 2018 from cogongrass individuals in north sites and south sites, where C- and E-type occur next to each other (1–20 m). The seeds from each seed parent were germinated at 30/20°C. The genotype of each ramet and each seedling was evaluated using the CAPS marker in the ITS region described above.

### Investigation of flowering phenology

Flowering periods of cogongrass were investigated in spring and fall in the natural habitats of north and south sites during 2016–2018 (Figure S2). In north sites, a total of 1,092 ramets were investigated in spring (July 2017 and May–June 2018) and 1,467 ramets in fall (November– December 2016 and October 2017) from 58 populations. In south sites, a total of 251 ramets were investigated in spring (April–June 2017) and 210 ramets in fall (November–December 2016 and October 2017) from six populations. Each population consisted of 2 to 92 ramets. Ramets in the flowering and non-flowering phases were sampled at >1-m intervals. The genotype of each ramet was evaluated using the CAPS marker in the ITS region described above.

Flowering phenology was monitored for C-, E-type, and artificially crossed F_1_ plants cultivated from 1 April to 30 November 2018 at the experimental farm of Kyoto University. The experimental farm was located between two hybrid zone, north and south sites. The survey was conducted for 141 accessions from 122 populations of the C-type and 83 accessions from 54 populations of the E-type and artificially generated F_1_ hybrids. The age of these plants were not controlled. The artificial F_1_s were generated by hand pollination between two ecotypes collected from the same prefecture, namely populations from Miyagi, Ibaraki, Ishikawa, Fukui, Shizuoka, Aichi, Osaka, and Wakayama, in 2010 (Miyoshi and Tominaga 2017) and 2017. Success of hybridisation was evaluated using the CAPS marker in the ITS region. Two hundred twenty-one individuals in which hybridisation was successful were used for the observation of flowering phenology. The plants were grown in plastic pots (15.9-cm diameter × 19.0-cm height) or clay pots (21.8-cm diameter × 17.5-cm height) containing paddy soil. The date on which the top of the first panicle emerged from a leaf sheath was defined as the flowering date.

### Investigation of seed sets

Together with the survey of flowering phenology in natural habitats in 2016–2018, panicles in the seed dispersal phase were collected in May to July for the C-type (21 populations) and E-type (23 populations) and in October to December for F_1_ (25 populations) in 2016–2018 (Figure S1). One hundred twenty-two panicles of C-type and 183 panicles of E-type were collected in spring and 228 panicles of F_1_ in fall. Seed set was evaluated by counting the spikelets (flowers) and seeds of each panicle. The genotype of each panicle was evaluated using the CAPS marker in the ITS region described above.

Hand pollinations were conducted between different accessions from the same ecotype: C-type × C-type, E-type × E-type, and F_1_ × F_1_. The crossing accession pairs were isolated in a mesh-covered cubic frame and their panicles were rubbed when their anthers and stigmas were matured. The seed sets of hand-pollinated C-type, E-type, and F_1_ plants in the experimental farm were evaluated (Table S3). Two accessions of the C-type were from south sites. Two accessions of the E-type were from north sites, and two from south sites. Four accessions of F_1_ were from north sites, and four from south sites. Pairs consisted of accessions from the same region and each accession was collected from different populations. These plants were crossed within a pair during April–May 2017 for the C- and E-types and during September–October 2018 for F_1_. Panicles were collected during May–June 2017 for the C- and E-types and during November–December 2018 for F_1_. The spikelets and seeds of each panicle were counted.

### Germinability tests

In total, 1055, 958, and 99 seeds from 16 populations of the C-type, 22 populations of the E-type, and 14 populations of F_1_, respectively, were subjected to germination test. The seeds were collected during the survey of seed sets in 2016–2018 (Figure S1). Seeds of cogongrass are non-dormant, and seedling emergence occurs soon after seed dispersal (Matumura et al. 1983; Shilling et al. 1997); therefore, seed germination tests were conducted immediately (within 2 months) after seed collection.

Under the outside condition, 271 seeds of the C-type and 433 seeds of the E-type were sown on 23 May and 13 June 2017 and on 20 June 2018. Fifty-eight seeds set on F_1_ were sown on 19 December 2016 and 4 December 2017. Germination tests were conducted on 4 December 2017 using 177 seeds of the C-type, as seeds set on F_1_ were too few to check the germinability of those sown in winter. The C-type seeds were collected during June–July 2017 in north and south sites and were stored at 4°C in a refrigerator. Seeds were sown on the surface of potting soil (Takii & Co., Ltd., Kyoto, Japan) in a plastic pot (23.2-cm diameter × 29.7-cm height) in the experimental farm of Kyoto University. Seed germination was observed daily for 1 month. For seeds set on F_1_, seed germination was observed daily until the following June.

Under controlled conditions, 784 seeds of the C-type and 525 seeds of the E-type were sown on 24 May, 16 June, and 21 August 2017 and on 20 June 2018 in a growth chamber (Biorton NC-220S, NK system, Japan). Forty-one seeds set on F_1_ were sown on 19 December 2016 and 4 December 2017. In addition to F_1_, germination tests were conducted on 4 December 2017 using 160 seeds of the C-type collected during June–July 2017 in north and south sites and stored at 4°C in a refrigerator. Seed germination tests were conducted on filter paper in Petri dishes under 12 h light/dark and 30/20°C, the optimal conditions for seed germination of cogongrass (Matumura et al. 1983; Mizuguti et al. 2002). In addition to natural seeds, seed germination tests were conducted using seeds produced by artificial crossing of the above-mentioned four pairs of F_1_ plants (Table S3). Seed germination tests were conducted on filter paper in Petri dishes under 12 h light/dark at 30/20°C, 25/15°C, 20/10°C, and 15/5°C using 103, 90, 90, and 109 seeds, respectively. Seed germination was observed daily for 1 month.

### Statistical analysis

All tests were two-tailed, and the significance level was set at *P* < 0.05. Hybridisation percentage (hybridised: 1/no: 0), flowering percentage (flowered: 1/no: 0), seed set percentage (set: 1/no: 0), and germination percentage (germinated: 1/no: 0) in the natural habitats were compared among genotypes (ecotypes and F_1_ hybrids) using generalised linear mixed models (GLMM) constructed using the package ‘lme4’ (Bates et al. 2015) in R 3.5.1 software (R Core Team 2018). The model assumed a binominal distribution and a logit link function. The genotypes were treated as fixed effects. The origin of the populations was treated as a random effect. Pairwise significant differences were identified by *post hoc* tests for GLMM (Tukey honestly significant difference test) using the package ‘multcomp’ (Hothorn et al. 2008) in R 3.5.1.

## Results

### Population genetic structure of cogongrass

To investigate the genetic structure of cogongrass in Japan, we developed 12 nuclear and one CAPS chloroplast markers (Figure S3; Table S4). Fifty-three SNPs were detected using the 12 nuclear markers from 223 accessions of cogongrass. Haplotypes in four regions, *ppc-C4*, ITS, EST72, and EST104, were not shared by the C- and E-types in Japan (Figure S3; Table S4), supporting the previously suggested genetic isolation of the two ecotypes (Mizuguti et al. 2004; Nomura et al. 2015). F_2_ genotypes were investigated using four molecular markers that can distinguish ecotypes and hybrids: ppc-C4, EST72, ITS_CAPS, and EST104_CAPS (Table S2). These four regions showed very low coefficients of linkage disequilibrium in an analysis of 28 putative F_2_ s, suggesting that they are located on different chromosomes (Table S5).

PCoA based on the 12 nuclear markers separated 223 accessions into three clusters along PCo1 (Figure 3B). The chloroplast CAPS marker, which is used to distinguish maternal ecotypes, revealed that the clusters at both sides of PCo1 corresponded to the C- and E-types (Figure 3B). The cluster in the middle of PCo1 was considered putative hybrids between ecotypes. In a STRUCTURE analysis, the number of optimum clusters was estimated to be *K* = 2 by Evanno *ΔK* (Figure 3C), implying that Japanese cogongrass populations are composed of two genetic clusters. All of the putative hybrids showed *q* values of around 0.5, suggesting that they had half of the genomes of the C-type and half of the E-type. A NEWPHYBRIDS analysis assigned the two ecotypes and putative hybrids to parental classes and the F_1_ hybrid class, respectively (Figure 3C). There was no putative F_2_ hybrid or backcross individual, except for one accession that belonged to the E-type cluster but carried the C-type haplotype of cpDNA. In the first survey of 223 populations using 12 nuclear markers, F_1_s were found in a few geographic locations, namely the north and south sites. We further investigated 127 accessions of F_1_ populations using the ITS CAPS marker, which was developed to discriminate the C-type, E-type, and F_1_. The distribution pattern of F_1_s was similar to the first survey. These F_1_ plants were found where C- and E-types share habitats. The majority (85%) of the F_1_ hybrids exhibited the E-type cpDNA haplotype, indicating that the E-type tends to be the maternal parent of hybrids, as supported by the hybridisation percentage in natural habitats (Figure 3D).

**Figure 3.**
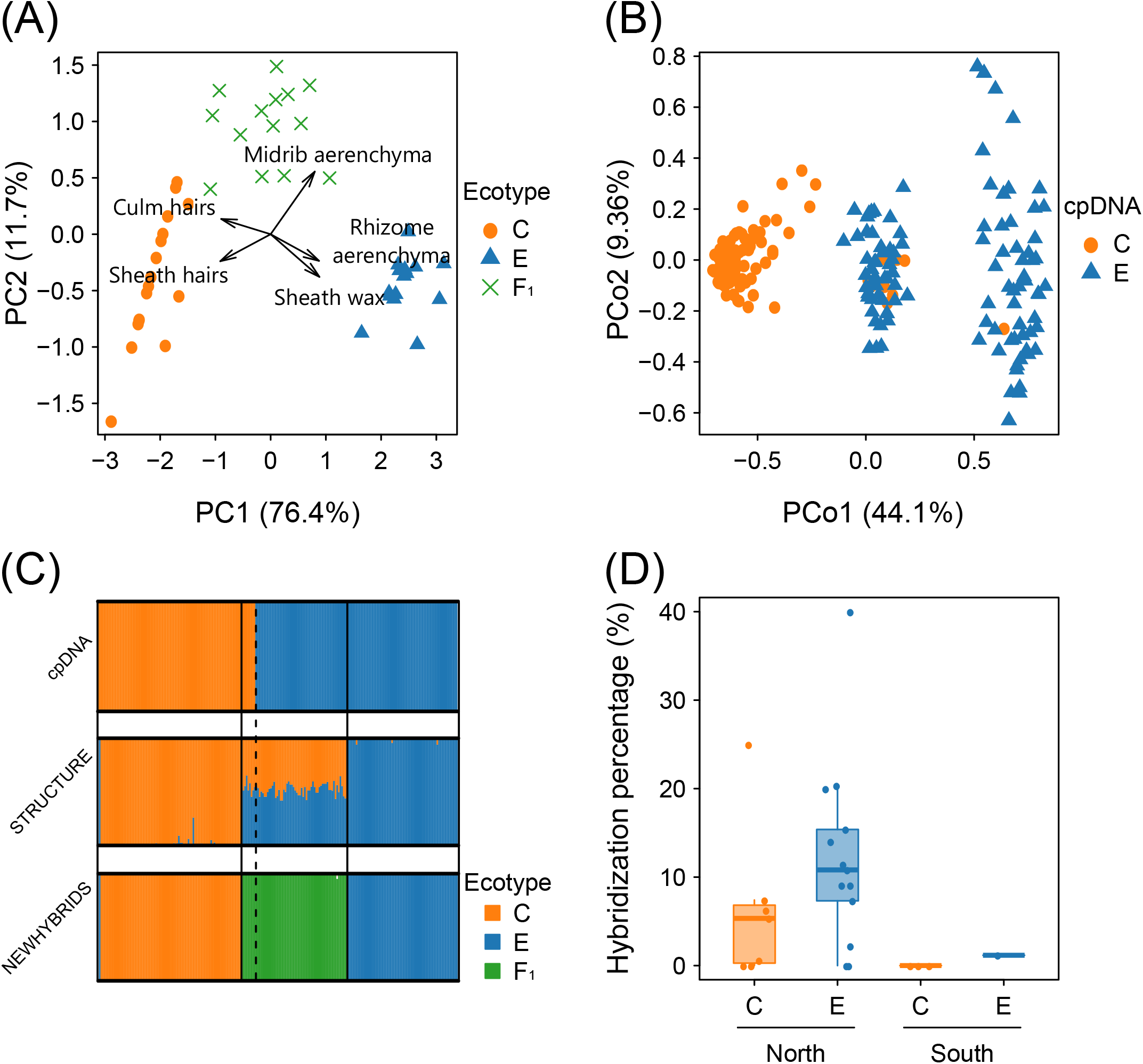
Differentiation between C- and E-type and hybridisation percentage. Results of PCA for morphological traits (A). Results of PCoA for 12 markers in nuclear DNA (B). Results of analysis for cpDNA and STRUCTURE and NEWHYBRIDS for 12 markers in nuclear DNA (C). Top, middle and bottom figures are results of analysis for cpDNA, STRUCTURE and NEWHYBRIDS. Hybridisation percentage in natural habitats (D). X-axis means genotypes of panicles (C- or E-type) and study sites (north and south sites). Points means hybridisation percentage per each population (surveyed panicles in north sites: C-type, n = 377; E-type, n = 395; south sites: C-type, n = 222; E-type, n = 86). Hybridisation percentage of E-type is significantly higher than that of C-type (GLMM, *P* < 0.01).

The occurrence and direction of hybridisation events between the C- and E-types were investigated in natural habitats sympatrically colonised by the C- and E-types (north and south sites). Five hundred ninety-nine seeds set on 50 panicles of the C-type, and 481 seeds set on 50 panicles of the E-type were collected from 10 and 14 populations in natural habitats, respectively. CAPS analysis of the ITS region of the progeny revealed a low percentage (~1.2%) of hybridisation in south sites (Figure 3D). In contrast, north site populations exhibited a higher percentage of hybridisation (~40%). The direction of hybridisation was asymmetric in both regions: significantly more F_1_ plants were observed among the seedlings from E-type seed parents (GLMM, *P* < 0.01). This result is consistent with the observation that the majority of F_1_ hybrids in the wild exhibit E-type cpDNA haplotypes (Figure 3C).

### Morphological traits of the ecotypes and F_1_

Culms, leaf sheathes (Tominaga et al. 1989; Mizuguti et al. 2003), and midrib aerenchyma (A. Nishiwaki & A. Mizuguti unpublished data) are different between the C- and E-types. The C-type had hairs on culm nodes and leaf sheathes and no wax on leaf sheaths, whereas the E-type had no hairs on culm nodes or leaf sheathes and had wax on the leaf sheaths (Figure 1, S4). The C-type also had a smaller ratio of midrib aerenchyma to midrib diameter than the E-type had. In addition to the previously identified ecotype-specific characteristics, we investigated rhizome aerenchyma based on the assumption that the habitat of soil moisture contents of the ecotypes may be associated with their rhizome structures. We found that the size of rhizome aerenchyma differed between ecotypes; the C-type had small aerenchyma, and the E-type had large aerenchyma.

F_1_ showed a wide range of variation in morphology; various numbers of hairs on culm, no or few hairs and little wax on sheathes, and an intermediate ratio of aerenchyma. As a result, F_1_ was located in an intermediated position between parental ecotypes on axis 1 of PCA (Figure 3A). The C- and E-types and F_1_ were divided into three groups by PCA. The axis 1 was associated with hairs on culms, sheath was, sheath hairs, and rhizome aerenchyma, indicating that they are the most diagnostic of hybridisation. Therefore, the C- and E-types and F_1_ were distinct, not only genetically but also morphologically.

### Flowering phenology and reproduction of F_1_

Population genetic analyses revealed that hybrid individuals were F_1_s, suggesting reproductive isolation of F_1_ plants from their parents (backcross) and prevention of F_1_ × F_1_ crossing. We first investigated the flowering period of the C- and E-types and F_1_ in the fields to assess the occurrence of reproductive isolation between F_1_ and the two ecotypes. The number of ramets at the flowering stage was counted in 64 populations. The genotype of each ramet was later investigated using the CAPS marker in the ITS region. Three hundred one (66.9%) of 450 ramets of the C-type and 345 (77.9%) of 443 ramets of the E-type were in the flowering stage in spring, in agreement with previous reports (Matumura and Yukimura 1980; Tominaga et al. 1989) (Figure 4A) (Tukey HSD, *P* < 0.001). In contrast, only 11 (2.0%) of 552 ramets of F_1_ flowered in spring. In fall, up to 48.0% of F_1_ ramets (581/1211) flowered, compared to only 5.3% of C-type and 1.6% of E-type ramets (Tukey HSD, *P* < 0.001). The flowering phenology of F_1_ plants was not affected by cpDNA type (Figure S5).

**Figure 4.**
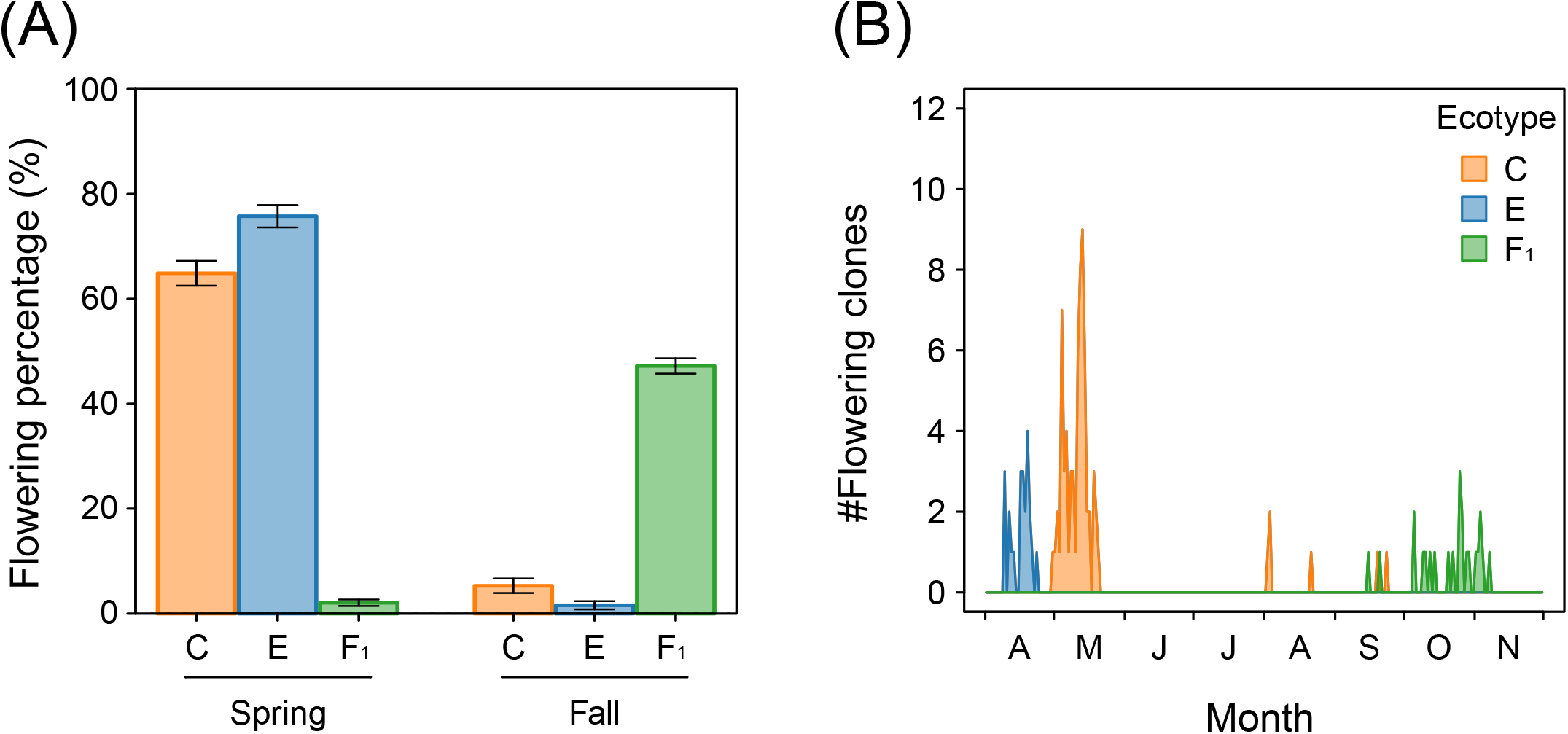
Flowering phenology of C-, E-type and F1. Flowering phenology in natural populations (A). Flowering percentage means (flowering ramets/all ramets) × 100 (surveyed ramets in spring: C-type, n = 407; E-type, n = 404; F1, n = 532; fall: C-type, n = 264; E- type, n = 252; F1, n = 1161). Values and error bars represent means ± s.e.m. Flowering percentage of F1 is significantly lower than that of C-type and E-type in spring while that of F1 is significantly higher than that of C- type and E-type in fall (Tukey HSD, *P* < 0.001). Flowering phenology in Kyoto experimental farm (B).

To validate the late flowering of F_1_ in natural habitats, we generated artificial F_1_ plants of eight reciprocal sets and monitored their flowering phenology from April to November in 2018 at the experimental farm. During mid-April, flowering of E-type accessions peaked (Figure 4B). Approximately 1 month later, flowering of the C-type peaked. During this period, none of the F_1_ plants flowered. In contrast, F_1_ sporadically flowered from September to November, as observed for wild F_1_ populations. The results indicate that hybridisation between the C- and E-types of cogongrass delays the flowering of F_1_ progeny to fall, which prevent F_1_s from crossing with the parental ecotypes.

Next, to clarify the mechanism of prevention of F_1_ × F_1_ crossing, we surveyed seed sets on F_1_ plants in natural habitats from 25 populations in fall. The C-type produced more flowers than the E-type (359.1 and 246.1 on average, respectively), as reported previously (Matumura et al. 1983). F_1_ plants produced 368.6 flowers on average, suggesting that they had the same ability to produce flowers as the C-type. The average seed set, however, was 0.12% (0.0–2.5%); thus, most of the F_1_ panicles in natural habitats did not carry mature seeds (Figure 5A). In contrast, our spring survey showed that the average seed set of the C- and E-types in natural habitats was approximately 70-fold higher than that of F_1_ plants (Tukey HSD, *P* < 0.001): 7.1% (0.0–70.6%) for the C-type and 9.0% (0.0–60.7%) for the E-type. A similar seed set percentage, 8.7% (0.6–24.7%), was observed in F_1_ plants when different populations were artificially crossed (Figure 5A), suggesting that F_1_ plants have an extremely low seed set in natural habitats, although they can produce offspring at rates similar to the C- and E-types.

**Figure 5.**
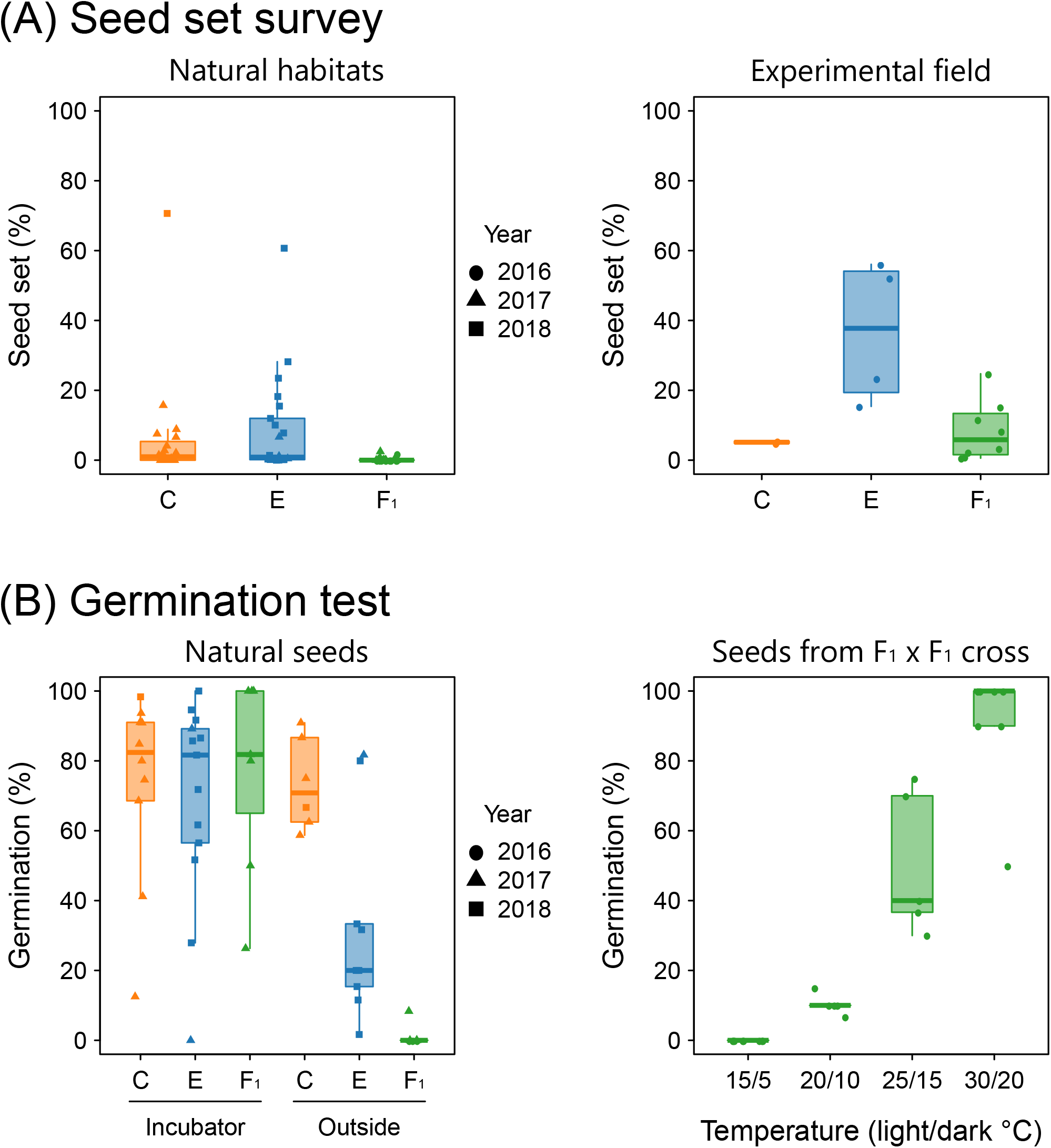
Viability in sexual reproduction of C-, E-type and F1. Seed set analysis in natural habitats and an experimental field (A). Symbols mean a year in which an experiment was conducted in natural habitats (surveyed panicles: C-type, n = 122; E-type, n = 183; F1, n = 228). Seed setting of F1 is significantly lower than that of C-type and E-type (Tukey HSD, *P* < 0.001). Seed set percentage of hand-pollinated ecotypes and F1 was measured in an experimental field of Kyoto University (surveyed panicles: C-type, n = 7; E-type, n = 9; F1, n = 20). Seed setting of F1 is not significantly differ from that of C-type (Tukey HSD, *P* = 0.996) and significantly lower than that of E-type (Tukey HSD, *P* < 0.01). Germination tests of seeds from natural habitats or artificial pollination (B). Left panel shows germination test of natural seeds under an incubator and an outside condition. Germination tests in incubator were conducted under 30/20°C, light/dark condition. C- and E-type seeds were sown in summer (May-August), while seeds set on F1 were sown in winter (December) (surveyed seeds under incubator: C-type, n = 784; E-type, n = 525; F1, n = 41; outside: C- type, n = 271; E-type, n = 443; F1, n = 58). Germination percentage of F1 is not significantly differ from that of E-type (Tukey HSD, *P* = 0.64) and significantly lower than that of C-type (Tukey HSD, *P* < 0.05) under the incubator condition. On the other hands, germination percentage of F1 is significantly lower than that of C-type (Tukey HSD, *P* < 0.001) and E-type (Tukey HSD, *P* < 0.05) under the outside condition. Right panel shows germination percentage of seeds produced from F1 × F1 cross under four temperature conditions (surveyed seeds: 5–15°C, n = 109; 10–20°C, n = 90; 15–25°C, n = 90; 20–30°C, n = 103). All points in three panels means seed set or germination percentage per each population or cross treatment.

We also investigated the viability of seeds derived from F_1_ × F_1_ crosses. Seeds collected from artificial F_1_ × F_1_ crosses were subjected to germination tests at various temperatures. As reported for the C- and E-types (Matumura et al. 1983; Mizuguti et al. 2002), the frequency of germination was high at 30/20°C, although none of the seeds germinated at 15/5°C (Figure 5B). Under the optimal conditions, mature seeds on F_1_ plants in natural habitats were also tested for germination immediately after seed collection, as seeds of cogongrass do not exhibit dormancy (Matumura et al. 1983; Shilling et al. 1997). A significantly lower and similar germination percentage compared to seeds of C- and E-type plants from natural habitats were observed in F_1_ plants, respectively (C-type, Tukey HSD, *P* < 0.05; E-type, Tukey HSD, *P* = 0.64) (Figure 5B). When the seeds were sown outside, however, the pattern of germination differed between F_1_ and the parental ecotypes (C-type, Tukey HSD, *P* < 0.001; E-type, Tukey HSD, *P* < 0.05). Seeds of C- and E-types that were sown in May to June had a higher germination percentage: 64.9% (176 seeds) for the C-type and 40.4% (179 seeds) for the E-type. In contrast, seeds from F_1_ plants in the natural habitats exhibited a low germination percentage when sown in December. Only 1 of the 58 seeds used for the experiment emerged by the following June. That seed germinated in April, when the average temperature was 16.4°C. We also used C-type seeds for the overwinter experiment due to the limited number of seeds from F_1_ plants. In total, 160 C-type seeds from six panicles (likely from different ramets), collected in June to July and stored at 4°C, were sown in the following December. Consistent with the result of seeds from F_1_ plants, a low germination percentage (7 of 177 seeds) was observed in C-type seeds sown in winter compared to in spring–summer: 0% germination in 117 seeds from five panicles and 11.7% germination in 60 seeds from one panicle (Figure S6) (GLMM, *P* < 0.001).

## Discussion

### Drastic shift in flowering phenology shapes population structures

In the genetic population structure of cogongrass in Japan, we found that hybrids between C- and E-type comprised only F_1_ generation except for one accession (Figure 3C). Four or five markers are enough to distinguish F_1_ from F_2_ or backcross individuals (Boecklen and Howard 1997). Therefore, used markers are a few in this study but classification by NEWHYBRIDS is appropriate. Origin of one accession with mismatched nuclear and chloroplast DNA types is unclear. It may be a chloroplast capture or an incomplete lineage sorting (Rieseberg and Soltis 1991; Kuritzin et al. 2016). The fact that F_1_ plants were identified in materials collected 30 years ago indicates that backcrossing and F_1_ × F_1_ crossing events have been prevented for at least 30 years. Investigation of F_1_ phenology revealed that they flowered in fall, whereas the parental ecotypes flowered in spring (Figure 4), which acts as a reproductive barrier to backcrossing. Our survey of the seed setting of F_1_ plants in natural habitats in fall suggests that the success rate of F_1_ × F_1_ crossing is 70-fold lower than that of the C- and E-types in spring (Figure 5A). Also, the germination percentage of the seeds set on F_1_ under the outside condition was markedly lower than that of the C- and E-types (Figure 5B). Notably, F_1_ plants possessed the potential for seed propagation, as did the C- and E-types. A high level of seed setting by F_1_ × F_1_ crossing was observed in hand pollination between plants from distinct populations (Figure 5A); the germination percentage was higher under warmer conditions (Figure 5B). Thus, the lower seed set and germination under natural conditions were likely caused by interactions with environmental parameters, such as temperature and population structure, which F_1_ plants confront as a result of drastic shifts in flowering phenology. Previous study demonstrated that some accessions of artificial F_1_ showed higher performance for biomass production than parental ecotypes under dry and wet conditions (Miyoshi and Tominaga 2017). Together, our data indicate that the peculiar F_1_ population structure of cogongrass is caused by loss of seed propagation, mediated by the drastic shift in flowering phenology, and it has been maintained by vigorous clonal reproduction via rhizomes (Figure 6).

**Figure 6.**
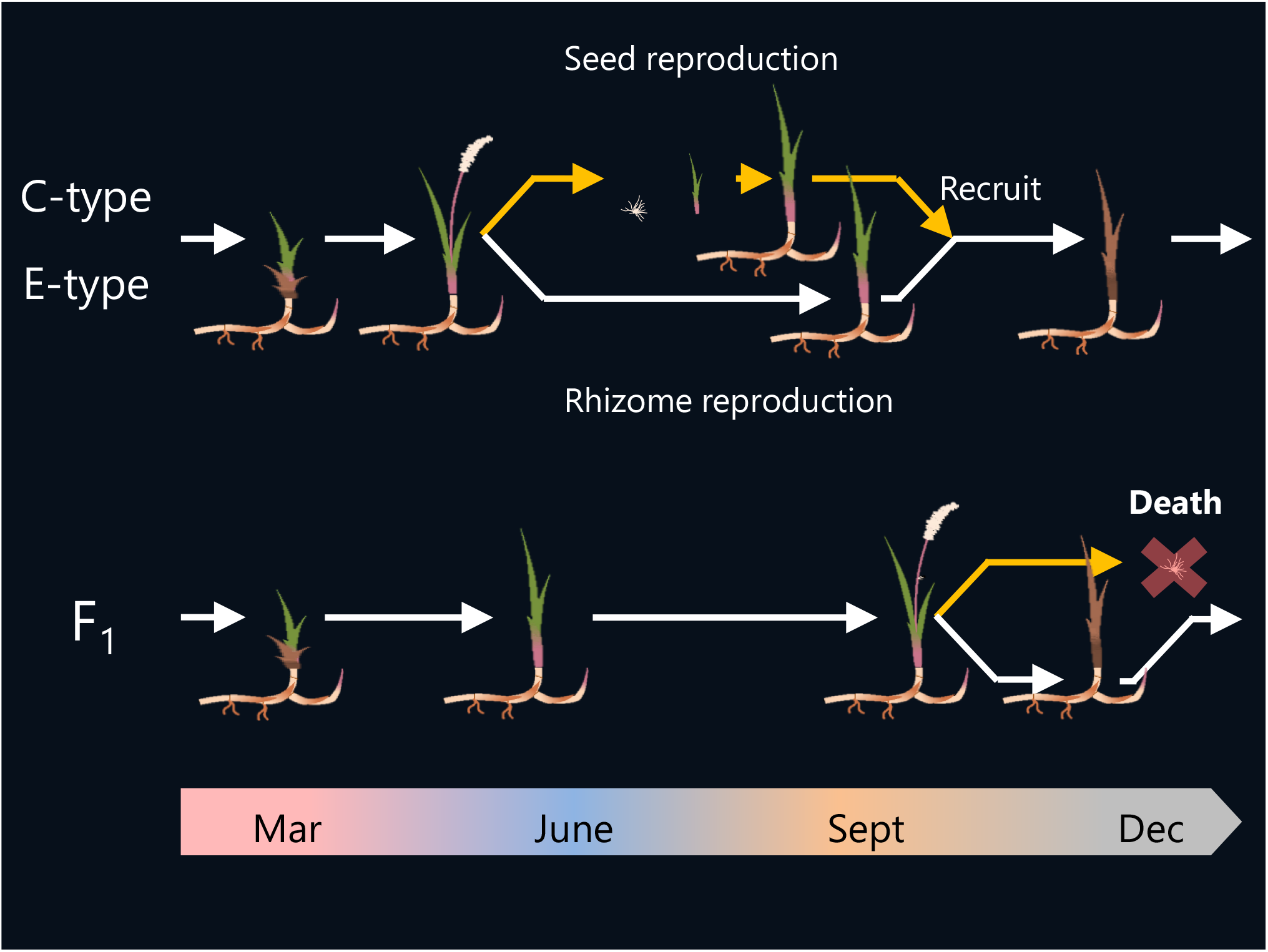
Mechanism of F_1_ domination in hybrid population. C- and E-type flower in spring and disperse their seeds in summer. Seedlings are likely to be establish, because seeds of C- and E-type are dispersed in a warm season. In contrast, F_1_ flowers in fall and disperse their seeds in winter. This suggests that seeds set on F_1_ are not able to germinate and are dead, because they are dispersed in a cold season.

Normal growth of artificially generated pseudo- F_2_ individuals (data not shown) indicates the existence of extrinsic mechanism(s) of excluding F_2_ plants in nature. Our data suggest that two events likely explain the absence of F_2_ plants: the exceptionally low seed-setting percentage of F_1_ plants and the low germination percentage of seeds dispersed in winter. The former mechanism(s) under natural conditions is unknown but is unlikely to be an environmental parameter such as temperature, based on the higher seed-setting percentage in our artificial crossing experiment in fall (Figure 5B) and the normal germination of F_1_ pollen in fall (T. Tominaga, unpublished data). The following are more plausible hypotheses: (1) Genetic diversity within/among F_1_ populations may affect seed setting percentage. Cogongrass can have a diverse seed setting percentages, which, due to its self-incompatible and outcrossing nature, would depend on the population genetic diversity (Shilling et al. 1997). F_1_ populations may hold insufficient genetic diversity to have a higher seed-setting percentage. (2) Non-synchronous flowering of F_1_ plants may affect the seed setting percentage. Contrary to the C- and E-types, flowering of F_1_ individuals occurred sporadically, and the flowering period was roughly 2 months, twofold longer than that of the C- and E-types (Figure 4B). Synchronisation of flowering would have a marked impact on the seed setting percentage of cogongrass. Further studies of the genetic population structures within/among F_1_ populations and the flowering period under natural conditions are needed to assess the low seed-setting percentage of F_1_ plants.

The exceptionally low germination percentage of the seeds dispersed in winter would also prevent the existence of F_2_ plants within hybrid populations. The germination percentage of the seeds sown in winter was 30-fold lower than that of seeds sown in spring. Given the low seed-setting percentage of F_1_ plants, roughly 0.05% that of the C- and E-types, an extremely low percentage of seedling establishment would be expected. In addition, our data may overestimate the germination percentage. In this study, we sowed the field-collected seeds in December, when cogongrass germination is suppressed by low temperature. In nature, however, seeds set and dispersed in early fall (such as September to early October) may germinate before winter. The seedlings are unlikely to overwinter because of insufficient rhizome development due to low temperature. Further investigation of seed dynamics, especially in nature, is required to understand the mechanism that prevents successful F_1_ × F_1_ crossing.

### Role of instant reproductive isolation on evolution

We identified a mechanism of instant reproductive isolation, in which almost complete reproductive isolation is established in one generation due to the self-incompatible and perennial nature of cogongrass. The former prevents generation of F_2_ plants, and the latter allowed the late-flowering hybrids to remain in the habitat by clonal reproduction (Figure 6). Furthermore, these characteristics have led to the population structure of hybrids with only F_1_ plants (Milne and Abbott 2008). Similarly, hybrid populations comprising only the F_1_ generation have been reported in perennial plants including herbaceous plants and woods (Nason et al. 1992; Kuehn et al. 1999; Milne et al. 2003; Kameyama et al. 2008; Milne and Abbott 2008; Zha et al. 2010; Nagano et al. 2015). These previous studies have proposed two major hypotheses on the absence of F_2_ and backcross hybrids: one is an extrinsic factor and the other is an intrinsic factor (Nason et al. 1992; Kuehn et al. 1999; Milne et al. 2003; Kameyama et al. 2008; Milne and Abbott 2008; Kameyama and Kudo 2011). The extrinsic factor hypothesis explains that a parental environment is unsuitable for F_2_ and backcross hybrids (Nason et al. 1992; Milne et al. 2003; Milne and Abbott 2008), while the intrinsic factor hypothesis explains that hybrid breakdown occurs (Kuehn et al. 1999; Kameyama et al. 2008; Kameyama and Kudo 2011). However, no direct factor determining population structure has been identified in above mentioned studies.

F_1_ progeny showed an intermediate morphology between the C- and E-types (Figure 3A, S4). The ecological roles of the traits investigated in this study are unknown, even in the parental ecotypes. However, the difference in morphology, particularly in rhizome aerenchyma, among the C-type, E-type, and F_1_ may influence the preference of habitat. In our observations, F_1_ plants were found in a wider range of soil moisture conditions than their parents (Nomura et al., in prep.), whereas the C- and E-types preferred relatively dry and wet habitats, respectively. This species is a worldwide noxious weed and has a wide range of distribution (Holm et al. 1977; MacDonald 2004; Burrell et al. 2015). This species is intentionally introduced as an ornament or forage grass and escapes unintentionally to natural habitats, expanding its distribution (MacDonald 2004; Cseke and Talley 2012; Burrell et al. 2015). If intentional/unintentional introduction provide F_1_ plants a chance to be distributed to a warm region where F_1_ can sexually reproduce, F_1_ will evolve into a new invasive taxon without backcrossing with parental species. Further studies of the role of rhizome aerenchyma in environmental preference may shed further light on the distribution of cogongrass in Japan.

### Regulation of flowering phenology

To ensure reproductive success, plants must regulate flowering to synchronise with optimal environmental conditions for seed production. A large number of molecular players are involved in the regulation of flowering in plants (Hill and Li 2016), which allows plants to fine-tune the flowering period. These players often function in an additive manner (Martin and Willis 2007; Martin et al. 2007; Buckler et al. 2009); therefore, crossing between individuals with different flowering periods often results in an intermediate flowering period. In contrast, marked delay of flowering has been observed in F_1_ hybrids of particular ecotypes/lines of *Arabidopsis thaliana* (L.) Heynh. (Koornneef et al. 1994; Henderson and Dean 2004) and *Sorghum bicolor* (L.) Moench (Murphy et al. 2014; Yang et al. 2014). In both cases, the parental ecotypes/lines have a disrupted form of a floral repressor or its activator in the floral regulatory pathway, which lifts the repression of flowering. Hybridisation of the two ecotypes/lines restores the flowering repression pathway because the hybrids carry functional alleles of each gene. The genes involved in the delayed phenology differ between *A. thaliana* and *S. bicolor* although their basic relationship as a floral repressor and its activator is identical. The genes related to the delayed phenology of F_1_ in cogongrass are unknown. We speculate that similar molecular mechanisms as in *S. bicolor* are involved in the delayed phenology in cogongrass because the two species are evolutionarily similar—both are in the tribe Andropogoneae. Genomic and transcriptomic approaches will enhance our understanding of the dynamic flowering shift in cogongrass.

### Conclusion

A novel phenotype derived from hybridisation between two ecotypes has major effects on the population structure of cogongrass. The fact that hybrid populations consist of almost of all F_1_ plants implies that the F_1_ progeny of C- and E-type cogongrass lose their sexual reproduction system and only reproduce asexually. Namely, hybridisation of the two ecotypes altered not only the flowering phenology but also the reproductive strategy of cogongrass. Considering that similar delays in flowering phenology are observed in other plant lineages (*e*.*g*. *A. thaliana* and *S. bicolor*), it is reasonable to speculate that hybridisations between independently evolved ecotypes may cause a drastic shift of flowering phenology even in other plants. Our findings will facilitate investigation of the ecological role of hybridisation in flowering phenology shifts in other species. Also, the molecular basis of the flowering shift is an exciting challenge for the future.

## Acknowledgement

Partly supported by the Japan Society for the Promotion of Science (21380015) and ESPEC Foundation for Global Environment Research and Technology to T.T.

## Supporting information

Figure S1

Title: Sampling location of each survey

Legends: Sampling location of flowering phenology survey (A) and sampling location of samples used for seed set (B) and germination test (C). The left panel is spring survey and the right panel is fall survey. Symbols mean a year in which a survey was conducted.

Figure S2

Title: CAPS markers in nuclear DNA

Legends: Complete length of ITS in cogongrass and a recognition site (A). Arrows and a triangle mean primers and a recognition site of *Dde*I, respectively. Grey bars mean not-deposited sequences to Genbank. Fragment length polymorphisms of ITS (B). ITS regions were amplified using ITS CAPS primer set and PCR products were digested by *Dde*I. Complete length of EST104 in cogongrass and a recognition site (C). Arrows and a triangle mean primers and a recognition site of *Rsa*I, respectively. Grey bars mean non-deposited sequences to Genbank. Fragment length polymorphisms of ITS (D). EST104 were amplified using EST104 CAPS primer set and PCR products were digested by *Rsa*I.

Figure S3

Title: Estimated haplotypes in C-, E-type and F_1_ and SNP sites in each region

Legends: The IDs correspond to haplotype IDs in each region (Table S4). Filled color means haplotypes found in C-, E-type and F_1_.

Figure S4

Title: Morphological traits of both ecotype and F_1_

Legends: Culm hairs (A), wax on leaf sheaves (B), hairs on leaf sheaves (C), ratio of aerenchyma diameter in a rhizome pith to rhizome diameter (D) and ratio of aerenchyma diameter to leaf midrib diameter (E).

Figure S5

Title: CpDNA type of flowering ramets in natural habitats

Legends: Filled and hatched areas mean flowering and non-flowering ramets, respectively. Orange and blue colors mean cpDNA of C- and E-type, respectively.

Figure S6

Title: Germination percentage of C-type sown in December under an incubator condition and an outside condition

Legends: Germination tests in an incubator were conducted under 30/20°C, light/dark condition (surveyed seeds under incubator, n = 160; outside, n = 177). Germination percentage under the incubator condition is significantly higher than that under the outside condition (*P* < 0.001).

Table S1. Used accessions

Table S2 Primer sets

Table S3 Seed set of hand-pollinated panicles

Table S4. SNPs in haplotypes of each region

Table S5 Linkage disequilibrium *r*^2^ for four regions

The English in this document has been checked by at least two professional editors, both native speakers of English. For a certificate, please see: http://www.textcheck.com/certificate/LIBpy9 and http://www.textcheck.com/certificate/ZPfIvS

## Notes

**Declaration of Interests** The authors declare no competing interests.

### Competing Interest Statement

The authors have declared no competing interest.

## Reference

Anderson, E. C., and E. A. Thompson. 2002. A model-based method for identifying species hybrids using multilocus data. Genetics 160:1217–1229.

Bates, D., M. Mächler, B. Bolker, and S. Walker. 2015. Fitting linear mixed-effects models using lme4. J. Stat. Softw. 67:1–48.

Boecklen, W. J., and D. J. Howard. 1997. Genetic analysis of hybrid zones : Numbers of markers and power of resolution. Ecology 78:2611–2616.

Buckler, E. S., J. B. Holland, P. J. Bradbury, C. B. Acharya, P. J. Brown, C. Browne, E. Ersoz, S. Flint-Garcia, A. Garcia, J. C. Glaubitz, M. M. Goodman, C. Harjes, K. Guill, D. E. Kroon, S. Larsson, N. K. Lepak, H. Li, S. E. Mitchell, G. Pressoir, J. A. Peiffer, M. O. Rosas, T. R. Rocheford, M. C. Romay, S. Romero, S. Salvo, H. S. Villeda, H. Sofia da Silva, Q. Sun, F. Tian, N. Upadyayula, D. Ware, H. Yates, J. Yu, Z. Zhang, S. Kresovich, and M. D. McMullen. 2009. The genetic architecture of maize flowering time. Science 325:714–718.

Buerkle, C. A., R. J. Morris, M. A. Asmussen, and L. H. Rieseberg. 2000. The likelihood of homoploid hybrid speciation. Heredity 84:441–451.

Burrell, A. M., A. E. Pepper, G. Hodnett, J. A. Goolsby, W. A. Overholt, A. E. Racelis, R. Diaz, and P. E. Klein. 2015. Exploring origins, invasion history and genetic diversity of *Imperata cylindrica* (L.) P. Beauv. (Cogongrass) in the United States using genotyping by sequencing. Mol. Ecol. 24:2177–2193.

Coyne, J. A., and H. A. Orr. 2004. Speciation. Sinauer, Sunderland, MA U.S.A.

Cseke, L. J., and S. M. Talley. 2012. A PCR-based genotyping method to distinguish between wild-type and ornamental varieties of *Imperata cylindrica*. J. Vis. Exp. 60:e3265.

Duenez-Guzman, E. A., J. Mavárez, M. D. Vose, and S. Gavrilets. 2009. Case studies and mathematical models of ecological speciation. 4. hybrid speciation in butterflies in a jungle. Evolution 63:2611–2626.

Evanno, G., S. Regnaut, and J. Goudet. 2005. Detecting the number of clusters of individuals using the software STRUCTURE: a simulation study. Mol. Ecol. 14:2611–2620.

Grant, P. R., and B. R. Grant. 1994. Phenotypic and genetic effects of hybridization in Darwin’s finches. Evolution 48:297–316.

Henderson, I. R., and C. Dean. 2004. Control of *Arabidopsis* flowering: the chill before the bloom. Development 131:3829–3838.

Hill, C. B., and C. Li. 2016. Genetic architecture of flowering phenology in cereals and opportunities for crop improvement. Front. Plant Sci. 7:1906.

Holm, L. G., D. L. Plucknett, J. V. Pancho, and J. P. Herberger. 1977. *Imperata cylindrica* (L.) Beauv. POACEAE (also GRAMINEAE). GRASS FAMILY. Pp. 62–71 in L. G. Holm, D. L. Plucknett, J. V. Pancho, and J. P. Herberger, eds. The World’s Worst Weeds, Distribution and Biology. University Press of Hawaii, Honolulu, USA.

Hothorn, T., F. Bretz, and P. Westfall. 2008. Simultaneous inference in general parametric models. Biometrical J. 50:346–363.

Jakobsson, M., and N. A. Rosenberg. 2007. CLUMPP: a cluster matching and permutation program for dealing with label switching and multimodality in analysis of population structure. Bioinformatics 23:1801–1806.

Johansen-Morris, A. D., and R. G. Latta. 2006. Fitness consequences of hybridization between ecotypes of *Avena barbata*: hybrid breakdown, hybrid vigor, and transgressive segregation. Evolution 60:1585–1595.

Kagawa, K., and G. Takimoto. 2018. Hybridization can promote adaptive radiation by means of transgressive segregation. Ecol. Lett. 21:264–274.

Kameyama, Y., T. Kasagi, and G. Kudo. 2008. A hybrid zone dominated by fertile F1s of two alpine shrub species, *Phyllodoce caerulea* and *Phyllodoce aleutica*, along a snowmelt gradien t. J. Evol. Biol. 21:588–597.

Kameyama, Y., and G. Kudo. 2011. Clarification of the genetic component of hybrids between *Phyllodoce caerulea* and *Phyllodoce aleutica* (Ericaceae) in Hokkaido, northern Japan. Plant Species Biol. 26:93–98.

Koornneef, M., H. Blankestijn-de Vries, C. Hanhart, W. Soppe, and T. Peters. 1994. The phenotype of some late-flowering mutants is enhance by a locus on chromosome 5 that is not effetive in the Landsberg erecta wild-type. Plant J. 6:911–919.

Kuehn, M. M., J. E. Minor, and B. N. White. 1999. An examination of hybridization between the cattail species *Typha latifolia* and *Typha angustifolia* using random amplified polymorphic DNA and chloroplast DNA markers. Mol. Ecol. 8:1981–1990.

Kuritzin, A., T. Kischka, J. Schmitz, and G. Churakov. 2016. Incomplete lineage sorting and hybridization statistics for large-scale retroposon insertion data. PLoS Comput. Biol. 12:1–20.

Lamichhaney, S., F. Han, M. T. Webster, L. Andersson, B. R. Grant, and P. R. Grant. 2018. Rapid hybrid speciation in Darwin’s finches. Science 359:224–228.

MacDonald, G. E. 2004. Cogongrass (*Imperata cylindrica*) -Biology, Ecology, and Management. CRC. Crit. Rev. Plant Sci. 23:367–380.

Mallet, J. 2007. Hybrid speciation. Nature 446:279–283.

Marques, I., A. Jürgens, J. F. Aguilar, and G. N. Feliner. 2016. Convergent recruitment of new pollinators is triggered by independent hybridization events in *Narcissus*. New Phytol. 210:731–742.

Martin, N. H., A. C. Bouck, and M. L. Arnold. 2007. The genetic architecture of reproductive isolation in louisiana irises: flowering phenology. Genetics 175:1803–1812.

Martin, N. H., and J. H. Willis. 2007. Ecological divergence associated with mating system causes nearly complete reproductive isolation between sympatric *Mimulus* species. Evolution 61:68–82.

Matumura, M., and T. Yukimura. 1980. The comparative ecology of intraspecific variants of the Chigaya, *Imperata cylindrica* var. *koenigii* (Alang-alang). (1) Habitats of the common and early flowering types of the Chigaya based on the vegetation characteristics. Res. Bull. Fac. Coll. Agric. Gifu Univ. 43:233–248. [In Japanese with English abstract]

Matumura, M., T. Yukimura, and S. Shinoda. 1983. Fundamental studies on artificial propagation by seeding useful wild grasses in Japan IX. Seed fertility and germinability of the intraspecific two types of Chigaya (Alang-alang), *Imperata cylindrica* var. *koenigii*. J. Japanese Grassl. Sci. 28:395–404.

Mavárez, J., C. A. Salazar, E. Bermingham, C. Salcedo, C. D. Jiggins, and M. Linares. 2006. Speciation by hybridization in *Heliconius* butterflies. Nature 441:868–871.

Milne, R. I., and R. J. Abbott. 2008. Reproductive isolation among two interfertile *Rhododendron* species: low frequency of post-F1 hybrid genotypes in alpine hybrid zones. Mol. Ecol. 17:1108–1121.

Milne, R. I., S. Terzioglu, and R. J. Abbott. 2003. A hybrid zone dominated by fertile F1s: maintenance of species barriers in *Rhododendron*. Mol. Ecol. 12:2719–2729.

Miyoshi, I., and T. Tominaga. 2017. Growth of hybrids between the common and early ecotypes of *Imperata cylindrica*. Grassl. Sci. 63:128–131.

Mizuguti, A., A. Nishiwaki, and M. Oyamada. 2002. Differences of seed germination characters between two types of *Imperata cylindrica* (L.) Beauv. characterized by flowering phenology. Grassl. Sci. 48:216–220. [In Japanese with English abstract]

Mizuguti, A., A. Nishiwaki, and Y. Sugimoto. 2004. Genetic difference between two types of *Imperata cylindrica* (L.) Beauv. characterized by flowering phenology. Grassl. Sci. 50:9–14.

Mizuguti, A., A. Nishiwaki, and Y. Sugimoto. 2003. Morphological differences between two types of *Imperata cylindrica* (L.) BEAUV. characterized by flowering phenology. Grassl. Sci. 49:324–329. [In Japanese with English abstract]

Murphy, R. L., D. T. Morishige, J. A. Brady, W. L. Rooney, S. Yang, P. E. Klein, and J. E. Mullet. 2014. *Ghd7* (*Ma6*) represses Sorghum flowering in long days: alleles enhance biomass accumulation and grain production. Plant Genome 7:1–10.

Murray, M. G., and W. F. Thompson. 1980. Rapid isolation of high molecular weight plant DNA. Nucleic Acids Res. 8:4321–4326.

Nagano, Y., A. S. Hirao, and T. Itino. 2015. Genetic structure of a hybrid zone between two violets, *Viola rossii* Hemsl. and *V. bissetii* Maxim.: dominance of F1 individuals in a narrow contact range. Plant Species Biol. 30:237–243.

Nason, J. D., N. C. Ellstrand, and M. L. Arnold. 1992. Patterns of hybridization and introgression in populations of oaks, manzanitas, and irises. A m. J. Bot. 79:101–111.

Nomura, Y., Y. Shimono, and T. Tominaga. 2015. Development of chloroplast DNA markers in Japanese *Imperata cylindrica*. Weed Res. 55:329–333.

Peakall, R., and P. E. Smouse. 2006. GENALEX 6: genetic analysis in Excel. Population genetic software for teaching and research. Mol. Ecol. Notes 6:288–295.

Peakall, R., and P. E. Smouse. 2012. GenALEx 6.5: genetic analysis in Excel. Population genetic software for teaching and research-an update. Bioinformatics 28:2537–2539.

Pritchard, J. K., M. Stephens, and P. Donnelly. 2000. Inference of population structure using multilocus genotype data. Genetics 155:945–959.

R Core Team. 2018. R: A language and environment for statistical computing. R Foundation for Statistical Computing, Vienna, Austria.

Rieseberg, L. H., and D. E. Soltis. 1991. Phylogenetic consequences of cytoplasmic gene flow in plants. Evol. Trends Plants 5:65–84.

Selz, O. M., R. Thommen, M. E. Maan, and O. Seehausen. 2014. Behavioural isolation may facilitate homoploid hybrid speciation in cichlid fis h. J. Evol. Biol. 27:275–289.

Shilling, D. G., T. A. Bewick, J. F. Gaffney, S. K. McDonald, C. A. Chase, and E. R. R. L. Johnson. 1997. Ecology, Physiology, and Management of Cogongrass (*Imperata Cylindrica*): Final Report. Florida Institute of Phosphate Research. Institute of Food and Agricultural Sciences, University of Florida, Gainesville, Florida.

Soltis, P. S., D. B. Marchant, Y. Van de Peer, and D. E. Soltis. 2015. Polyploidy and genome evolution in plants. Curr. Opin. Genet. Dev. 35:119–125.

Soltis, P. S., and D. E. Soltis. 2009. The role of hybridization in plant speciation. Annu. Rev. Plant Biol. 60:561–588.

Stephens, M., N. J. Smith, and P. Donnelly. 2001. A new statistical method for haplotype reconstruction from population data. A m. J. Hum. Genet. 68:978–989.

Taylor, S. J., L. D. Rojas, S. W. Ho, and N. H. Martin. 2013. Genomic collinearity and the genetic architecture of floral differences between the homoploid hybrid species *Iris nelsonii* and one of its progenitors, *Iris hexagona*. Heredity (Edinb). 110:63–70.

Tominaga, T., H. Kobayashi, and K. Ueki. 1989. Geographical variation of *Imperata cylindrica* (L.) Beauv. in Japan. J. Japanese Grassl. Sci. 35:164–171.

Tominaga, T., A. Nishiwaki, A. Mizuguti, and T. Ezaki. 2007. Weed Monograph 5. *Imperata cylindrica* (L.) Beauv. J. Weed Sci. Technol. 52:17–27 [In Japanese].

Wood, T. E., N. Takebayashi, M. S. Barker, I. Mayrose, P. B. Greenspoon, and L. H. Rieseberg. 2009. The frequency of polyploid speciation in vascular plants. Proc. Natl. Acad. Sci. 106:13875–13879.

Yang, S., R. L. Murphy, D. T. Morishige, P. E. Klein, W. L. Rooney, and J. E. Mullet. 2014. Sorghum phytochrome B inhibits flowering in long days by activating expression of *SbPRR37* and *SbGHD7*, repressors of *SbEHD1*, *SbCN8* and *SbCN12*. PLoS One 9:e105352.

Yasuda, K., and H. Shibayama. 2006. Primer sets for DNA amplification of the noncoding regions of the chloroplast genome in the grass family. J. Weed Sci. Technol. 51:146–151.

Zha, H. G., R. I. Milne, and H. Sun. 2010. Asymmetric hybridization in *Rhododendron agastum*: a hybrid taxon comprising mainly F1s in Yunnan, China. Ann. Bot. 105:89–100.

