## Supplementary material for "Drastic shift in flowering phenology, an instant reproductive isolation mechanism, explains the population structure of *Imperata cylindrica* in Japan": Figure S1

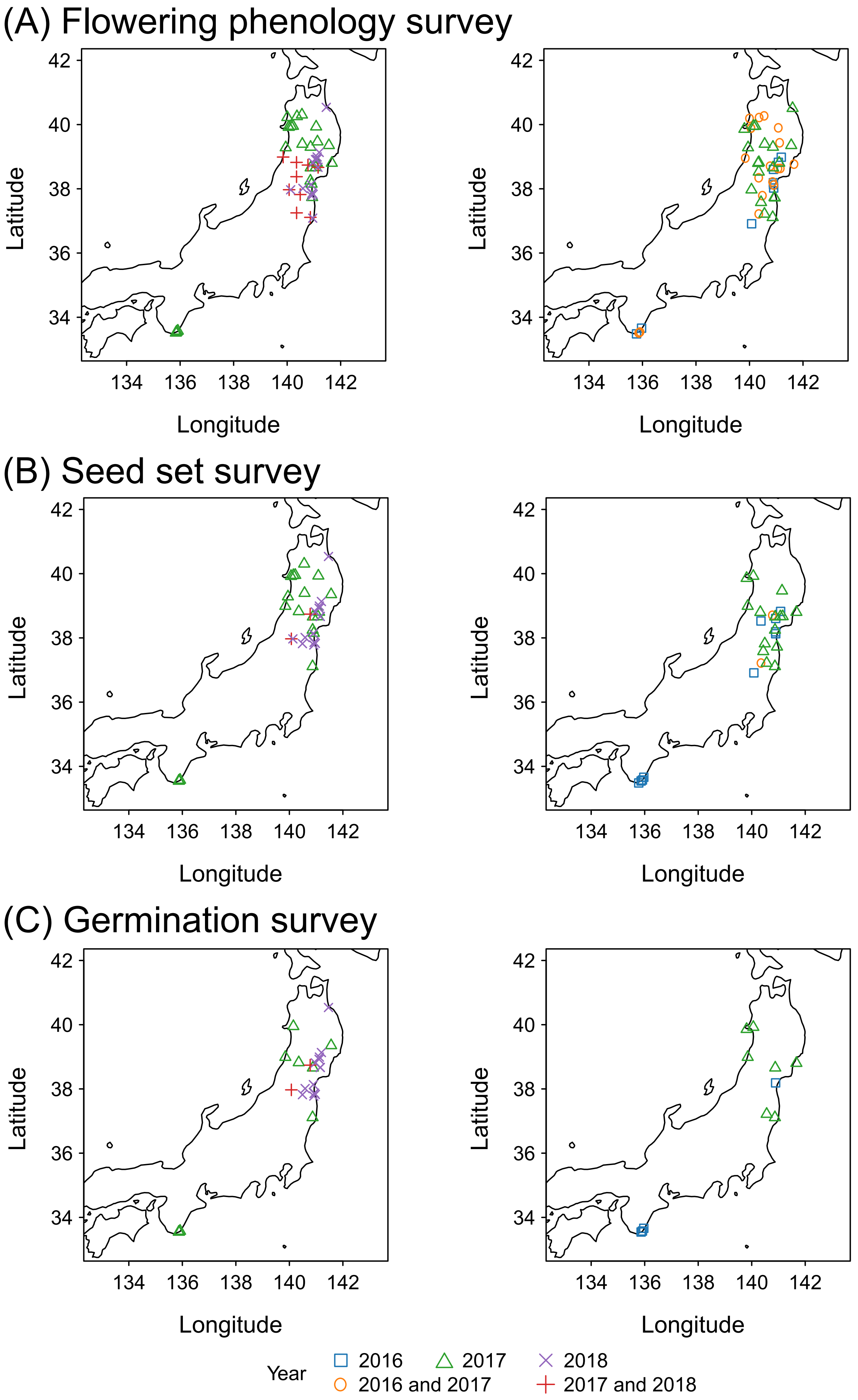


Figure S1

Title: Sampling location of each survey

Legends: Sampling location of flowering phenology survey (A) and sampling location of samples used for seed set (B) and germination test (C). The left panel is spring survey and the right panel is fall survey. Symbols mean a year in which a survey was conducted.
