## Supplementary material for "Drastic shift in flowering phenology, an instant reproductive isolation mechanism, explains the population structure of *Imperata cylindrica* in Japan": Figure S2

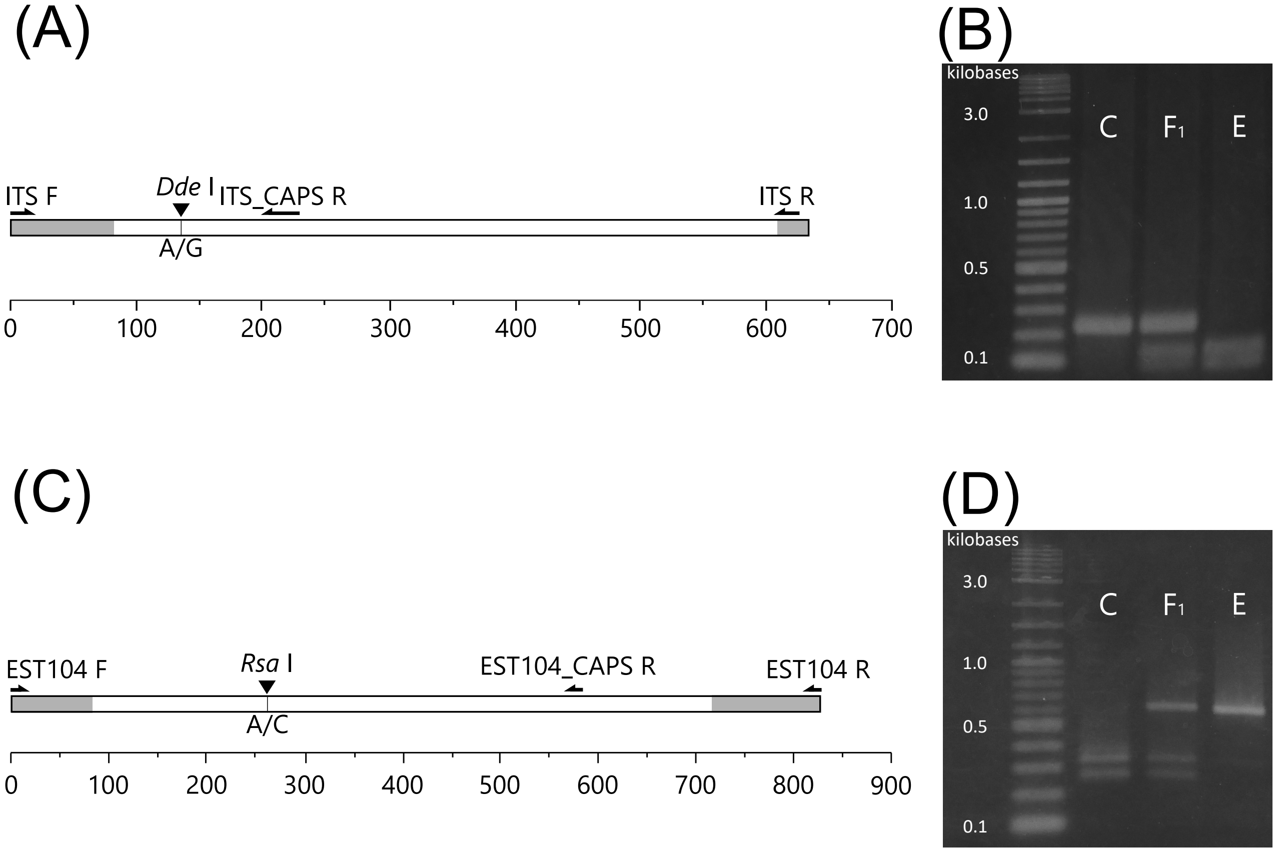


Figure S2

Title: CAPS markers in nuclear DNA

Legends: Complete length of ITS in cogongrass and a recognition site (A). Arrows and a triangle mean primers and a recognition site of *DdeI*, respectively. Grey bars mean not-deposited sequences to Genbank. Fragment length polymorphisms of ITS (B). ITS regions were amplified using ITS CAPS primer set and PCR products were digested by *DdeI*. Complete length of EST104 in cogongrass and a recognition site (C). Arrows and a triangle mean primers and a recognition site of *RsaI*, respectively. Grey bars mean non-deposited sequences to Genbank. Fragment length polymorphisms of ITS (D). EST104 were amplified using EST104 CAPS primer set and PCR products were digested by *RsaI*.
