## Supplementary material for "Drastic shift in flowering phenology, an instant reproductive isolation mechanism, explains the population structure of *Imperata cylindrica* in Japan": Figure S3

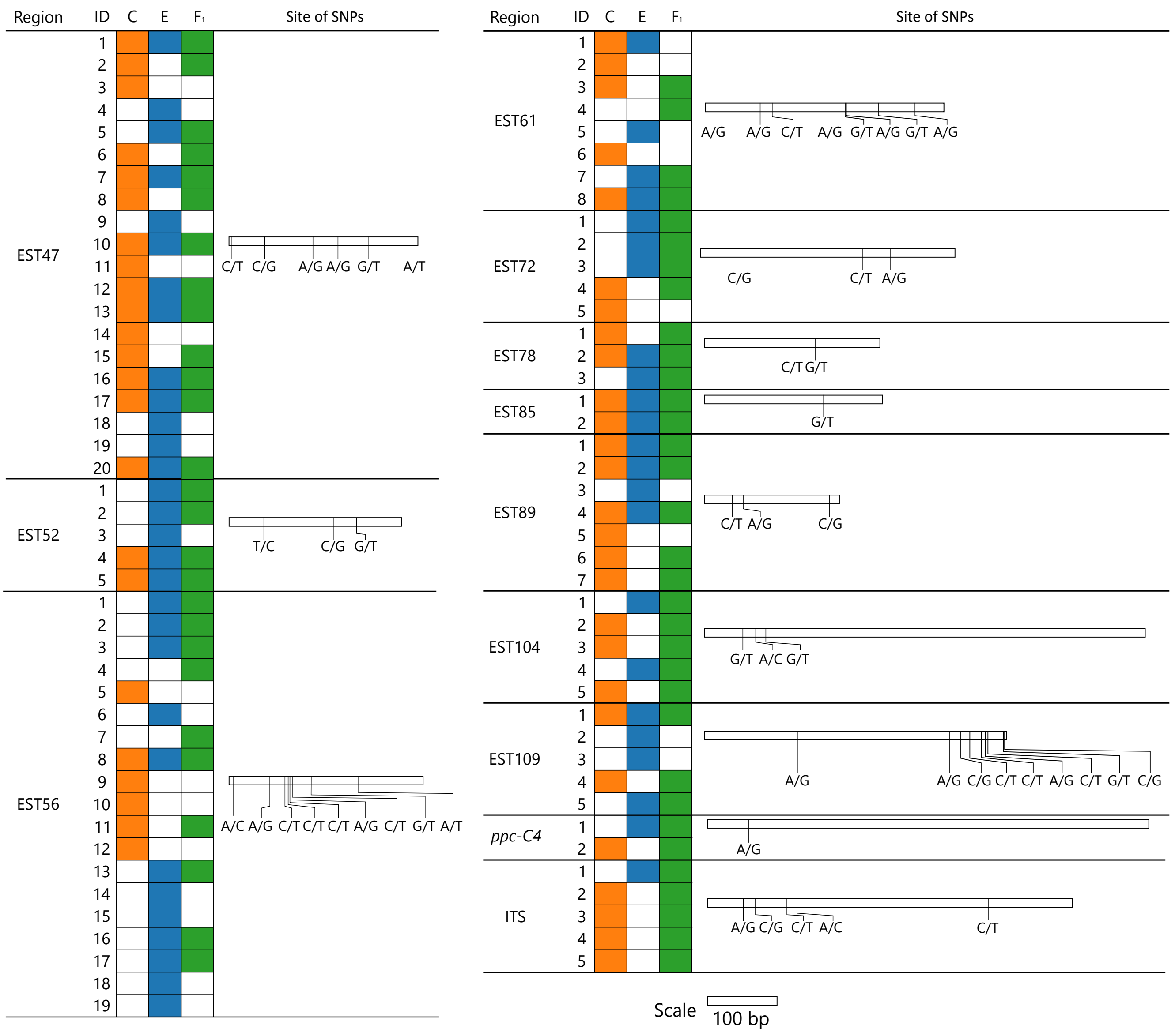


Figure S3

Title: Estimated haplotypes in C-, E-type and F1 and SNP sites in each region

Legends: The IDs correspond to haplotype IDs in each region (Table S4). Filled color means haplotypes found in C-, E-type and F1.
