## Supplementary material for "Drastic shift in flowering phenology, an instant reproductive isolation mechanism, explains the population structure of *Imperata cylindrica* in Japan": Figure S4

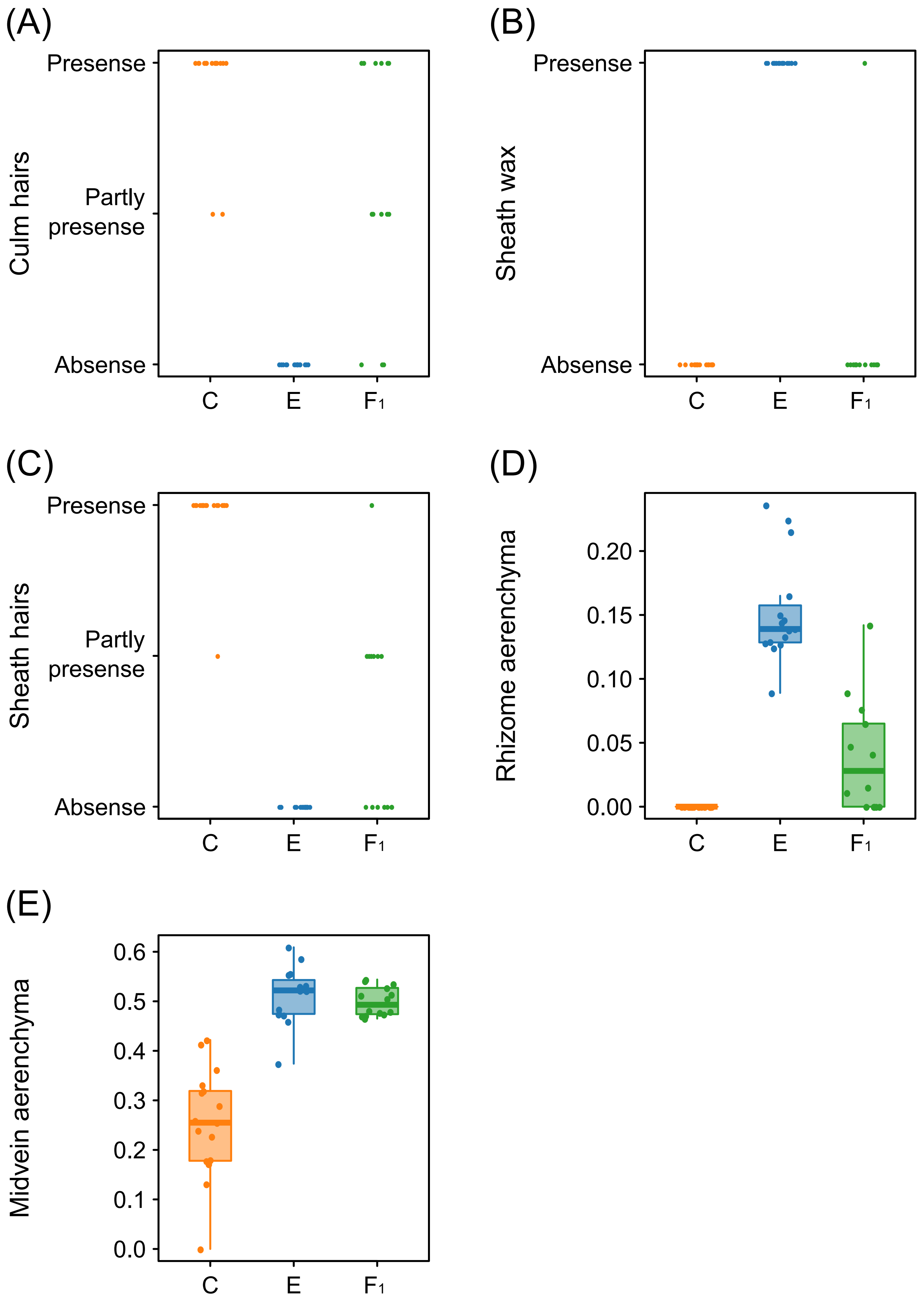


Figure S4

Title: Morphological traits of both ecotype and F1

Legends: Culm hairs (A), wax on leaf sheaves (B), hairs on leaf sheaves (C), ratio of aerenchyma diameter in a rhizome pith to rhizome diameter (D) and ratio of aerenchyma diameter to leaf midrib diameter (E).
