## Supplementary material for "Drastic shift in flowering phenology, an instant reproductive isolation mechanism, explains the population structure of *Imperata cylindrica* in Japan": Figure S5

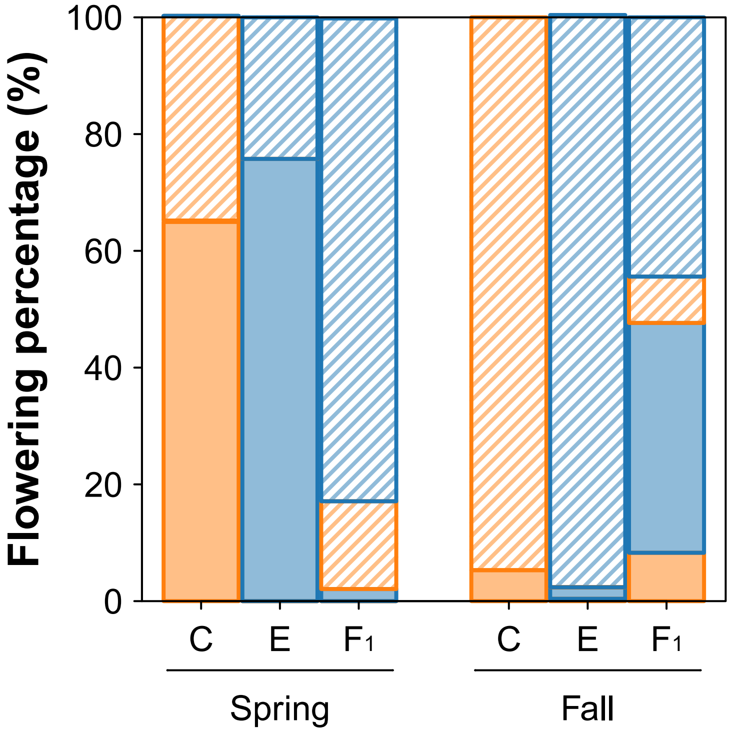


Figure S5

Title: CpDNA type of flowering ramets in natural habitats

Legends: Filled and hatched areas mean flowering and non-flowering ramets, respectively. Orange and blue colors mean cpDNA of C- and E-type, respectively.
