## Supplementary material for "Drastic shift in flowering phenology, an instant reproductive isolation mechanism, explains the population structure of *Imperata cylindrica* in Japan": Figure S6

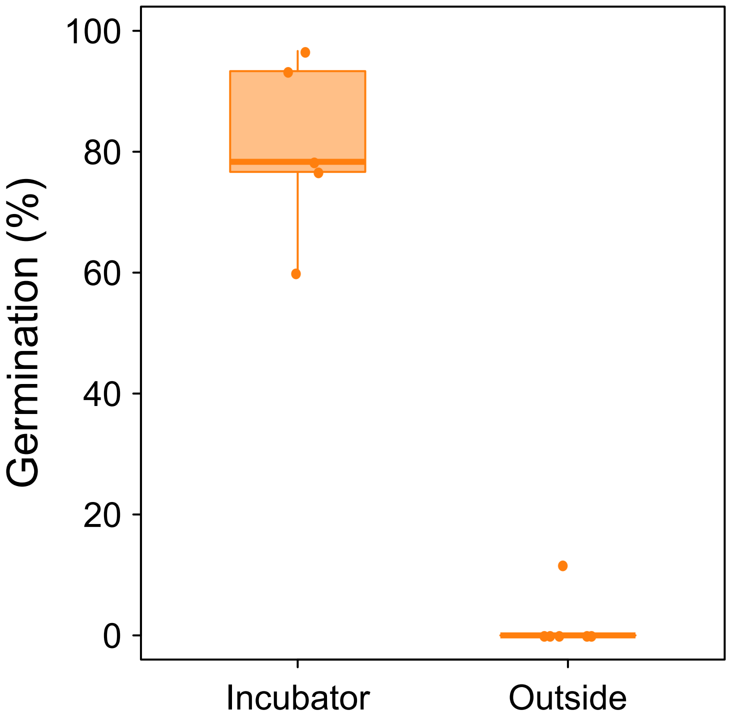


Figure S6
Title: Germination percentage of C-type sown in December under an incubator condition and an outside condition
Legends: Germination tests in an incubator were conducted under 30/20°C, light/dark condition (surveyed seeds under incubator, n = 160; outside, n = 177). Germination percentage under the incubator condition is significantly higher than that under the outside condition (*P* < 0.001).
